# Cerebral perfusion and metabolism coupling during a critical time window provides rapid assessment of cardiac arrest severity and prognosis in a preclinical model

**DOI:** 10.1101/785972

**Authors:** R. H. Wilson, C. Crouzet, M. Torabzadeh, A. Bazrafkan, N. Maki, J. Alcocer, B. J. Tromberg, B. Choi, Y. Akbari

## Abstract

Improved quantitative understanding of the dynamic relationship among cerebral blood flow, oxygen consumption, and electrical activity is important to clinicians treating acute brain injury. Such knowledge would elucidate the neurovascular response to ischemia, helping to potentially guide treatment. Using a multimodal optical imaging platform and a clinically-relevant rat model of cardiac arrest (CA) and cardiopulmonary resuscitation (CPR), we continuously measured cerebral blood flow (CBF), brain tissue oxygenation (StO_2_), cerebral metabolic rate of oxygen (CMRO_2_), and cerebral electrical activity (electrocorticography; ECoG). Multiple phases of cerebral hemodynamic recovery, with different degrees of mismatch between CBF and CMRO_2_, were observed following CPR. At 1 min post-resuscitation, we observed that the ratio CBF/CMRO_2_ is indicative of CA duration/severity and prognostic (with 87% accuracy) of short-term neurological recovery measured by the re-initiation of ECoG activity. These measurements provide the earliest known metrics for assessment of CA severity and prognosis post-CPR. Interestingly, the accuracy of this information is lost beyond 2-3 minutes post-CPR, highlighting a critical, easily overlooked, period immediately post-CPR. These metrics do not require pre-resuscitation data, underscoring translational potential in emergency-response settings when pre-CA information is unavailable. These metrics encourage validation in human studies, potentially offering real-time feedback during CA/CPR to optimize neurological outcome.

## Introduction

Cardiac arrest (CA) afflicts over 565,000 people annually in the United States (1). Survivors of CA typically sustain significant brain damage due to cerebral ischemia, reperfusion injury, and compromised cerebral autoregulation (2, 3). To improve patient outcomes, it is essential to better understand the complex response of the brain to CA and cardiopulmonary resuscitation (CPR). Specifically, it is crucial to quantitatively monitor the relationship between cerebral blood flow (CBF) and brain metabolism following CPR. After hypoxic-ischemic injury (e.g., CA), reperfusion (e.g., CPR) can deliver oxygenated blood to the brain to support energy production (4), but this massive influx of oxygen can also cause neuronal injury if it is not metabolized efficiently (5). As a result, relying on only perfusion data is insufficient to accurately assess the hemodynamic response of CA patients, and a complementary measure of cerebral metabolism is required for correct assessment of injury severity and prognostication of recovery (6). By monitoring CBF and brain metabolism in tandem, mismatches between these two quantities can be identified and corrected to improve patient outcome (7). This may be particularly critical during the initial minutes (or even seconds) post-CPR (8), when the most rapid hemodynamic changes occur (9, 10) and interventions targeting CBF might be most effective (11, 12).

Currently, hemodynamic status is typically monitored in emergency and intensive care settings by measuring blood pressure and blood gas concentration from the radial or femoral artery. However, these measurements are sometimes not continuously available during and immediately after CPR, occur distant from the brain, and are often not informative of cerebral hemodynamic processes (9). Methods for direct measurement of oxygen consumption in the brain are typically invasive (13), (14) and do not provide any direct information about perfusion. Non-invasive CBF measurement can be achieved with techniques such as MRI (15) and combined PET/CT (16), but those methods do not have the temporal resolution required to monitor rapid changes in cerebral perfusion and metabolism. Therefore, it remains difficult to obtain accurate real-time feedback on CBF and metabolism concurrently to guide emergent clinical intervention to optimize neurological outcome for CA patients.

To help address this unmet clinical need, optical spectroscopy and imaging have been recently used as both investigative and clinical tools (17–21). These methods employ visible-to- near-infrared (VIS-NIR) illumination onto the tissue and detection of the remitted light via elements such as a spectrometer, photomultiplier tube, or camera (21, 22). Here, within our previously-described (23–25) translational rat model of CA and CPR, we use high-speed multispectral spatial frequency domain imaging (SFDI) (24), laser speckle imaging (LSI) (23), and electrocorticography (ECoG) (23, 24, 26), to monitor CBF, brain oxygenation, cerebral metabolic rate of oxygen (CMRO_2_), and cerebral electrical activity. The ratio between CBF and CMRO_2_ quantifies the degree of mismatch between cerebral perfusion and metabolism, and serves as a metric of cerebral autoregulation (27). This ratio previously was employed in studies involving MRI (27, 28) and near-infrared spectroscopy (NIRS) (29) to assess the health of the brain in response to stimuli or hypoxic-ischemic events.

Here, in our preclinical model, we find that the CBF/CMRO_2_ ratio <u>*as early as 1 min post-CPR*</u> can accurately distinguish between shorter (less severe) CA and more prolonged CA. This result is of great potential importance for the clinical scenario in which a CA patient presents to first responders, emergency medicine physicians, or intensive care physicians who may lack knowledge of the exact time when CA occurred prior to achieving return of spontaneous circulation (ROSC). Additionally, the ratio of CBF and CMRO_2_ within 1 min from conclusion of CPR can predict (with 87% accuracy) the time that ECoG activity resumes. Previously, we showed that the area under the CBF curve over the first ~5 min post-CPR can predict ECoG burst time (9), but that technique required *a priori* knowledge of the total perfusion required for bursting to begin. By contrast, the perfusion and metabolism metrics reported here only require knowledge of CBF and CMRO_2_ in the first minute post-CPR. These metrics can help inform urgent clinical decision making in the critical period immediately post-CPR.

## Methods

### Animal Preparation

All procedures described in this protocol were approved by the Institutional Animal Care and Use Committee (IACUC) at the University of California, Irvine (protocol number 2013-3098-01) and conformed to ARRIVE (Animal Research: Reporting in Vivo Experiments) guidelines. Ten male Wistar rats (age 10-16 weeks, weight ~300-400 g), were used in this study. Based on the level of variability observed in our previous studies for optically-measured hemodynamic parameters (9, 10) (typical standard deviations of ~15-27% of mean), this number of rats was deemed sufficient for the hypothesis-generating pilot study described in this report. The animal preparation procedures are described in detail elsewhere (23, 24). Rats were fasted with three pellets the night before the experiment as standard procedure for our CA experiments. On the day of the experiment, rats were anesthetized with isoflurane and endotracheally intubated to enable controlled breathing with a ventilator. Then, epidural screw electrodes were implanted for ECoG, and a partial craniectomy (4 mm right-to-left x 6 mm anterior-to-posterior) was performed to expose a portion of the right sensory and visual cortex for optical imaging. Four epidural screw ECoG electrodes (one which served as the reference electrode) were implanted in the skull. Two of these electrodes were located toward the front of the brain (2 mm anterior to bregma, 2.5 mm lateral to bregma) over the motor cortices, one was located atop the visual cortex (5.5 mm posterior to bregma, 4 mm left of bregma), and the reference electrode was located in the posterior region of the brain (3 mm posterior to lambda), over the cerebellum. The femoral artery was cannulated to enable arterial blood gas sampling and blood pressure monitoring while the femoral vein was cannulated to enable intravenous drug delivery. The cranial window, ECoG electrodes, and optical imaging system (described below) are shown in Fig. 1(a).

**Fig. 1.**
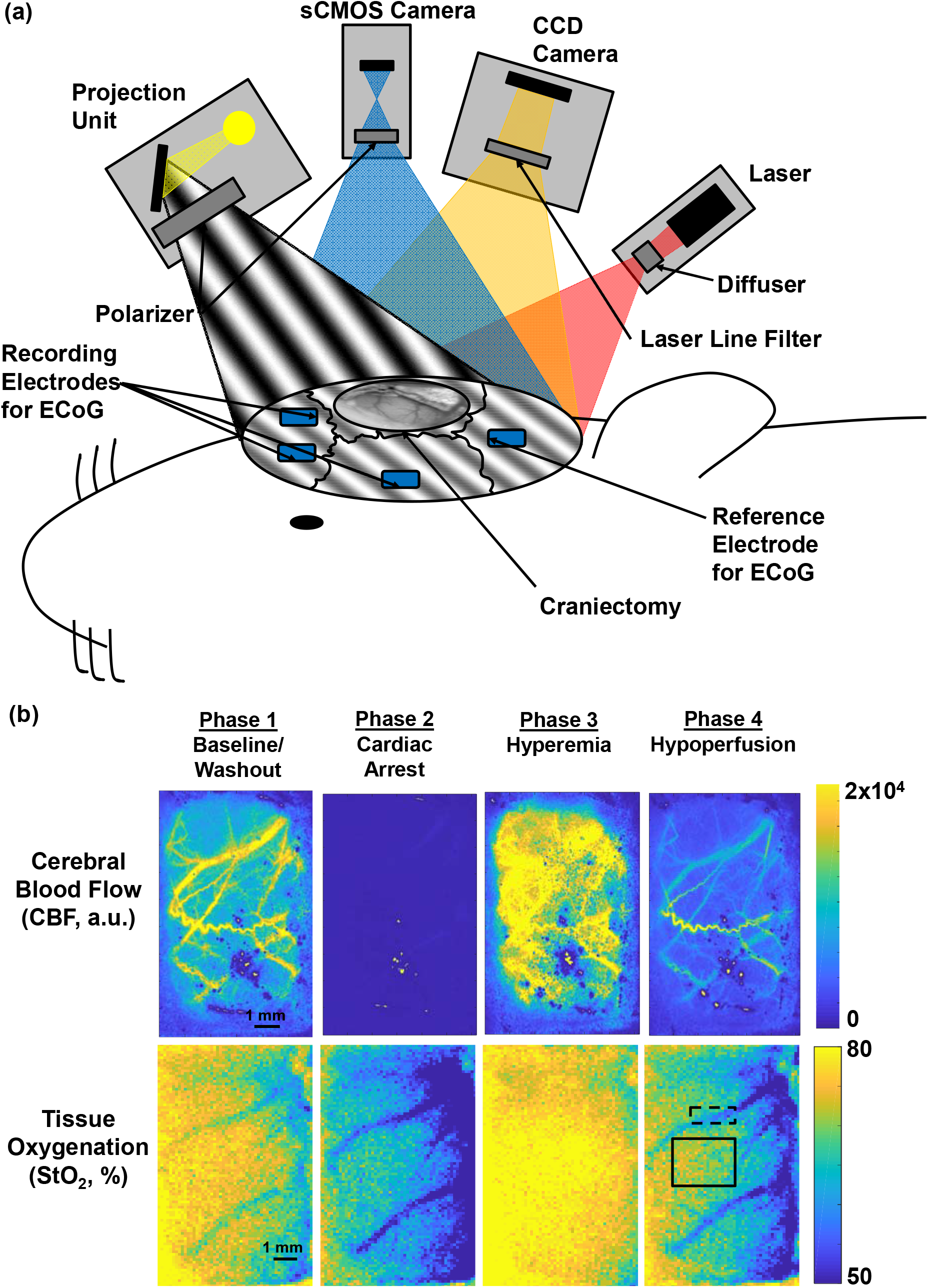
(a) Instrumentation for high-speed LSI and SFDI. Spatially-modulated LEDs and a sCMOS camera were used for SFDI. An 809 nm laser and 60 fps camera were used for LSI. The rapid LSI and SFDI system was integrated into an “animal intensive care unit” setup for monitoring the response of the brain to CA and CPR in a preclinical rat model. A craniectomy was performed to expose a ~6 mm x 4 mm region of the brain for imaging, and four EcOG electrodes were implanted for monitoring cerebral electrical activity. (b) Notable differences in CBF and oxygenation (StO_2_) were imaged during four phases: baseline, CA, hyperemic response to CPR, and post-hyperemic hypoperfusion. At baseline, normal cerebral perfusion and oxygenation were observed. During CA, blood flow was completely absent from the brain and cerebral oxygenation dropped rapidly, indicative of oxygen consumption. During post-ROSC hyperemia, states of hyper-perfusion and hyper-oxygenation were observed. During hypoperfusion, CBF stabilized at a level below baseline, yet increased metabolic activity was observed, reflected by a reduction in oxygenation.

### Laser Speckle Imaging (LSI)

The LSI system is described in detail in [7]. A long-coherence-length 809 nm laser (Ondax, Monrovia, CA) was the light source. A diffuser (ThorLabs, Inc., Newton, NJ) was mounted between the laser and the tissue to obtain near-uniform illumination on the exposed brain region. The remitted light was then isolated with a laser line filter and images were acquired at 60 Hz using a CCD camera (Point Grey Research Inc., Richmond, BC, Canada) with an exposure time T of 10 ms. For each region of interest (ROI) selected within the craniectomy, mean speckle flow index (SFI) values were calculated and used to create time-resolved curves. Relative SFI curves were calculated using a sliding median filter of 10 s in length. Baseline was defined as the mean SFI value calculated over the minute prior to asphyxia, resulting in a relative SFI value at baseline of 100. Unless otherwise specified, this pre-asphyxial time period was chosen as baseline due to post-anesthesia emergence and consequent cerebral hyperemia. However, in a scenario in which pre-CA data is unavailable, a different time period can be chosen as the baseline and changes relative to that new baseline can be assessed in the same manner. The SFI obtained using the above procedure was used as our measure of CBF in this report.

### Spatial Frequency Domain Imaging (SFDI)

The SFDI setup is described in detail in [8]. Briefly, we used three light emitting diodes (LEDs, 655 nm, 730 nm, 850 nm) as light sources that were coupled into a spatial light modulator to project spatial frequency patterns of the light onto the tissue. For detection of backscattered light, a scientific complementary metal-oxide semiconductor (sCMOS) camera (Hamamatsu Photonics) was employed. The acquisition sequence (DC projection, followed by a square-wave pattern at each of three spatial phases, repeated serially over all wavelengths) was repeated to achieve an effective frame rate of ~14 Hz. A two-step fitting procedure was incorporated to arrive at two-dimensional maps of reduced scattering coefficient (μs’), oxyhemoglobin concentration (ctHbO_2_), deoxyhemoglobin concentration (ctHb), total tissue hemoglobin concentration (ctHb_tot_), and tissue oxygen saturation StO_2_ = ctHbO_2_/ctHb_tot_.

### Relative cerebral metabolic rate of oxygen (rCMRO_2_) calculation

The baseline values (ctHb_o_, ctHb_tot,o_) and changes from baseline (ΔctHb, ΔctHbtot) in the deoxyhemoglobin and total hemoglobin concentrations from the SFDI measurements were combined with CBF values from the LSI measurements to calculate relative cerebral metabolic rate of oxygen (rCMRO_2_), using a previously-employed (30) equation:

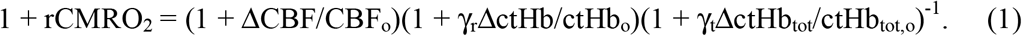

In Eq. (1), ΔCBF, ΔctHb, and ΔctHb_tot_ are the changes in CBF, deoxy-hemoglobin, and total hemoglobin, respectively, relative to their baseline values (CBF_0_, ctHb_0_, ctHbtot_0_). The constants γ_r_ and γ_t_ are related to the venous and arterial contributions to hemoglobin content, and were set to 1, which is within a range typically used for these parameters (30). As was the case for the CBF data, a pre-asphyxial time period was chosen as the baseline for the CMRO_2_ calculation unless otherwise specified.

### Definitions of flow-metabolism mismatch, coupling, and uncoupling

The flow-metabolism ratio CBF/CMRO_2_ was calculated at each time point by dividing the value of (1 + ΔCBF/CBF_o_) obtained with LSI by the value of (1 + rCMRO_2_) obtained with SFDI and LSI via Eq. (1) (29). This ratio was used to quantify *flow-metabolism mismatch*. Specifically, CBF/CMRO_2_ > 1 corresponded to a mismatch for which CBF exceeded metabolic demand, and CBF/CMRO_2_ < 1 represented a mismatch for which CBF was insufficient to meet metabolic demand. Separately, we defined periods of *flow-metabolism coupling* as time windows during which CBF and CMRO_2_ exhibited similar rates of change, and we defined periods of *flow-metabolism uncoupling* as time windows during which the slopes of CBF and CMRO_2_ had opposite signs, or where one slope was nonzero and the other was zero.

### Electrocorticography (ECoG)

Each screw electrode was connected to a PZ2 preamplifier (Tucker-Davis Technologies, Inc., Alachua, FL), which had a 0.35 Hz high-pass filter for detection of standard ECoG signals. A noise test (using the *pwelch* power spectral density function in MATLAB) was performed to ensure that there were no significant environmental sources that would reduce the signal-to-noise ratio for our measurements. Raw ECoG data were processed using custom MATLAB (Mathworks Inc., Natick, MA) code. DC bias was removed by de-trending the data. Noise and artifacts across channels were reduced with common average referencing (31). Then, a 60Hz notch filter and a 1-150 Hz bandpass filter were applied to the data. To lessen computational burden, signals were downsampled to 600Hz (23). ECoG burst frequency (defined as bursts/min) was used as a metric to quantify the extent of cerebral electrical recovery, as it has been shown to correlate with neurological outcome (32, 33).

### Cardiac Arrest (CA) and Cardiopulmonary Resuscitation (CPR)

At the beginning of the CA/CPR experiment, the isoflurane level was decreased from 2.0% to 0.5-1.0% and the inhaled gas mixture changed from 50% O_2_ + 50% N_2_ to 100% O_2_. After two min, isoflurane delivery was turned off and washed out by delivering room air (21% O_2_). This washout period is essential to mitigate effects of isoflurane on CBF and brain function. During this period, a neuromuscular blocking agent (1 mL of 2 mg/kg Vecuronium; 1 mL of heparinized saline) was administered intravenously to provide the ventilator with complete control of respiration. After three min, asphyxia was induced by turning off the ventilator for a fixed time period of 5 or 7 min, causing progressive hypoxic hypercarbic hypotension. For this pilot study, rats were not randomized into different CA duration groups, and investigators were not blinded to the CA duration or post-CA recovery. CA was identified when pulse pressure <10 mmHg and systolic pressure <30 mmHg. These conditions mimic pulseless electrical activity, a common type of CA (26).

Forty-five seconds prior to starting CPR, the ventilator was re-started with 100% oxygen given to the animal (respiratory rate = 85 breaths/min, PIP = 17.5-18.5 cmH2O, PEEP = 3 cmH2O at 2.5 LPM). Immediately before initiating CPR, intravenous administration of 0.01 mg/kg epinephrine, 1 mmol/kg sodium bicarbonate, and 2 mL of heparinized saline was performed. Then, CPR was conducted via external/closed chest compressions until return of spontaneous circulation (ROSC) was achieved (as determined by the blood pressure measured from the femoral artery). The duration of CPR was typically ~1 min. After ROSC, continuous monitoring (blood pressure, heart rate, ECoG, LSI, SFDI) of the animal was performed for ~1.5-2.0 hrs, followed by euthanasia with pentobarbital.

## Results

### Spatial mapping of cerebral perfusion and oxygen extraction in CA/CPR model

Fig. 1(b) shows representative maps of CBF and brain oxygenation during four distinct phases of the CA/CPR experiment. During Phase I (baseline/washout), CBF and oxygenation were constant until isoflurane washout began, at which point the CBF and CMRO_2_ both increased as the animal began to wake up. During Phase II (CA), asphyxia led to a rapid decrease in CBF and oxygenation due to progressive development of hypotension and eventual CA. During Phase III (hyperemia), immediately following CPR, a rapid increase in CBF and oxygenation occurred. During Phase IV (hypoperfusion), CBF stabilized at a level below baseline, but oxygen extraction increased, leading to a decrease in brain oxygenation. These trends are in agreement with our previously-reported results in this animal model (9, 10).

### Temporal dynamics of cerebral perfusion and metabolism in CA/CPR model

Figs. 2-3 illustrate the dynamic relationship between CBF and CMRO_2_ in a representative rat. During Phase I, CBF (Fig. 2(a); green), deoxy-hemoglobin (Fig. 2(a); blue), CMRO_2_ (Fig. 2(a); magenta), and ECoG activity (Fig. 2(b)) all increased during isoflurane washout. During Phase II, ECoG showed electrocerebral silence within ~30 sec following onset of asphyxia concomitantly with a decrease in systemic blood pressure (Fig. 2(c)). Also during Phase II, a large decrease in CBF, oxy-hemoglobin (Fig. 2(a); red), and CMRO_2_ was observed, with a large (~50%) increase in deoxy-hemoglobin, as oxygen extraction occurred in the absence of perfusion. During Phase III (Fig. 3), ROSC was associated with a hyperemic state, yet electrocerebral silence persisted. During Phase IV (Fig. 3), CBF decreased to a stabilized level, deoxy-hemoglobin increased, and ECoG activity resumed. These Phase IV dynamics signified increased oxygen extraction relative to perfusion, coinciding with increased neuronal activity.

**Fig. 2.**
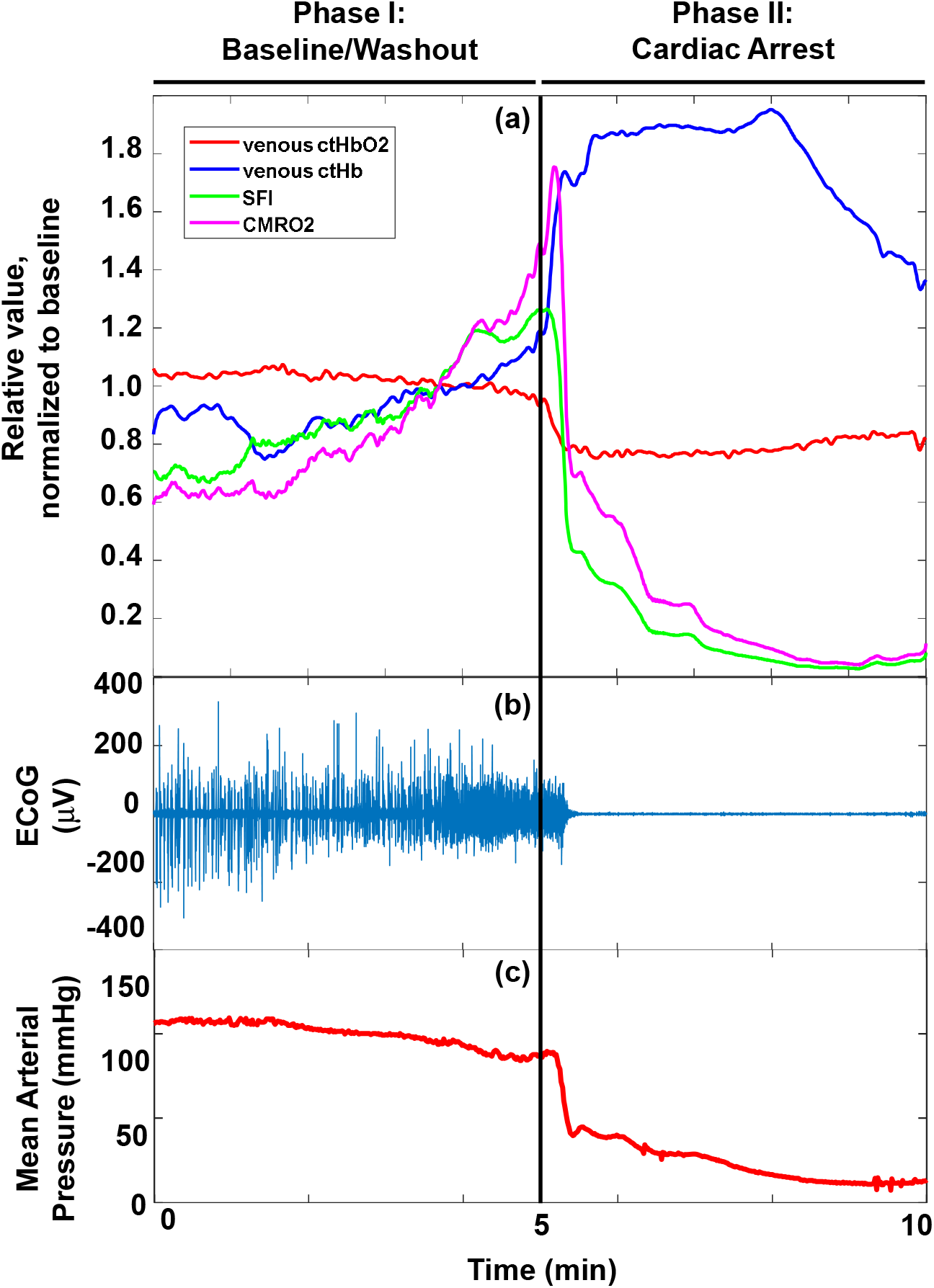
Different phases of CBF and metabolism were observed during isoflurane washout and subsequent onset of CA. (a) CBF (green), oxy-hemoglobin concentration (ctHbO_2_, red), deoxy-hemoglobin concentration (ctHb, blue), and CMRO_2_ (magenta), averaged over a region of interest for the same rat as in Fig. 1 during baseline (Phase I) and CA (Phase II). Note the rise in CMRO_2_ was slightly higher than the rise in CBF during washout, and this becomes much more dramatic during the initial seconds after asphyxia, suggesting flow/metabolism decoupling. (b) The ECoG signal was robust during Phase I, but reached pulseless electrical activity within ~30 sec after onset of asphyxia (Phase II). (c) Mean arterial pressure decreased sharply at the same time as onset of pulseless electrical activity.

**Fig. 3.**
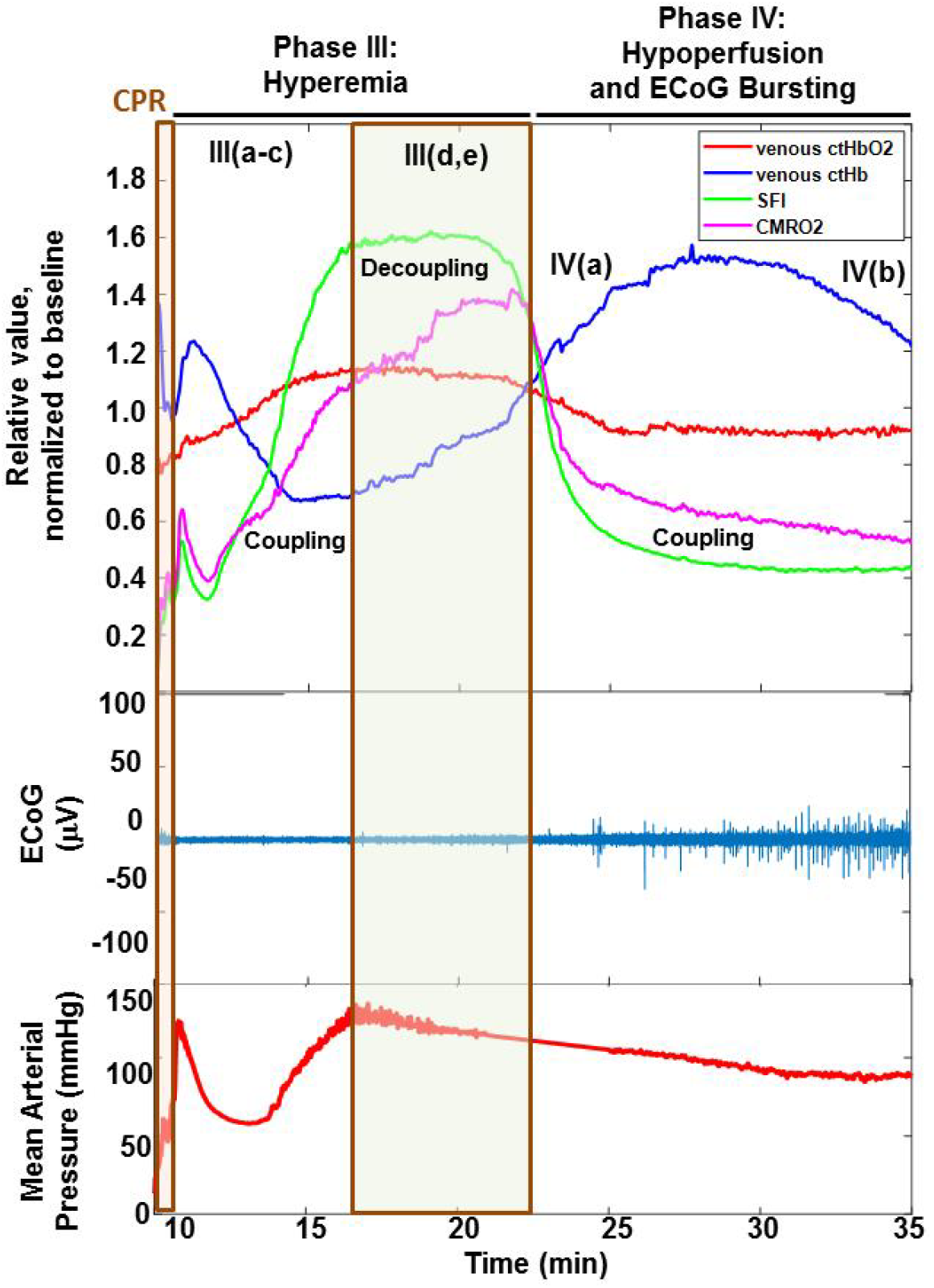
CBF (green), oxy-hemoglobin concentration (ctHbO_2_, red), deoxy-hemoglobin (ctHb, blue), and rCMRO_2_ (magenta), averaged over a region of interest for the same rat as in Fig. 2, are shown during CPR (orange shaded window), hyperemia (Phase III), and hypoperfusion (Phase IV). In addition, the associated whole-band ECoG signal and mean arterial pressure are shown. Soon after a decrease in CBF following hyperemia, ECoG bursting resumed and deoxy-hemoglobin increased, with oxygen extraction linked to the increased neuronal activity. Note that these hemodynamic changes observed during phases III and IV are not reflected in the waveform of mean arterial pressure, suggesting a decoupling between cerebral and peripheral hemodynamics. Multiple phases of cerebral flow-metabolism coupling and uncoupling occurred during Phase III. Flow-metabolism coupling (denoted as “coupling” in the figure) is defined as a period of concomitant changes in CBF and CMRO_2_ with similar slopes. Flow-metabolism uncoupling (denoted as “uncoupling” in the figure) is defined as a period during which CBF and CMRO_2_ have opposite slopes or one has a nonzero slope and the other has roughly zero slope.

We identified five sub-phases during Phase III (Fig. 3). During Phase III(a), which lasted for only ~1 min post-CPR, a transient increase in deoxy-hemoglobin, followed immediately by a transient decrease in CBF and CMRO_2_, was observed. During Phase III(b), CBF, oxyhemoglobin, and CMRO_2_ increased, and deoxy-hemoglobin decreased. During Phase III(c), CMRO_2_ and CBF continued to increase, but oxy-hemoglobin reached a plateau and deoxyhemoglobin began to increase, in a manner similar to that seen in Phase I. During Phase III(d), CMRO_2_ and deoxy-hemoglobin continued to increase, but oxy-hemoglobin slightly decreased while CBF reached a plateau. During Phase III(e), a noticeable transient decoupling between flow and metabolism was observed, as CMRO_2_ reached a plateau while CBF decreased sharply. This decoupling phase immediately preceded the initial ECoG burst. Six of the 10 rats exhibited either a period like Phase III(e) or a period during which CMRO_2_ increased only slightly during Phase III(c) while CBF was changing rapidly.

Phase IV (Fig. 3) contains two main sub-phases. During Phase IV(a), CBF and CMRO_2_ decrease sharply, oxy-hemoglobin continues to decrease gradually, and deoxy-hemoglobin increases. During Phase IV(b), CBF has stabilized at a level below pre-CA baseline and deoxy-hemoglobin gradually reaches a steady value. The end of hyperemia coincides with initial ECoG bursting and the transition between Phases III and IV, marked by the intersection of the CBF and CMRO_2_ curves. Following initial burst, ECoG recovery occurs, likely causing increased cerebral oxygen extraction that can cause the increase in deoxy-hemoglobin. This critical period of transition between Phases III and IV is evident from the combination of the CBF and oxygenation data but cannot be determined from the mean arterial pressure.

### Flow-metabolism coupling and uncoupling post-CPR may be influenced by CA duration

Fig. 4 shows CBF and CMRO_2_ for two representative rats: one with a 5 min asphyxial period and earlier time to initial ECoG burst frequency (top), and one with a 7 min asphyxial period and delayed recovery of burst frequency (bottom). For the rat with the shorter CA and earlier ECoG bursting, the CBF and CMRO_2_ dynamics are coupled throughout the reperfusion period. This similarity in the CBF and CMRO_2_ lineshapes, with the magnitude of the CBF change exceeding the magnitude of the CMRO_2_ change, is similar to that observed in stimulus-evoked CBF and CMRO_2_ measurements in healthy subjects (34). For the rat with the longer CA and delayed ECoG bursting, a longer period occurred before recovery of CBF and CMRO_2_, and the flow-metabolism dynamics were less coupled during hyperemia. Specifically, toward the middle of the hyperemic period (shaded box), CBF and CMRO_2_ trended in opposite directions, suggesting a mismatch between periods of increased blood flow and periods of higher metabolic demand in this rat. In four of the five rats with prolonged (7 min) asphyxia, notable differences were observed between the rate of change of CBF and the rate of change of CMRO_2_ during the reperfusion period. These periods of uncoupling were observed in only two of the five rats with shorter (5 min) asphyxia.

**Fig. 4.**
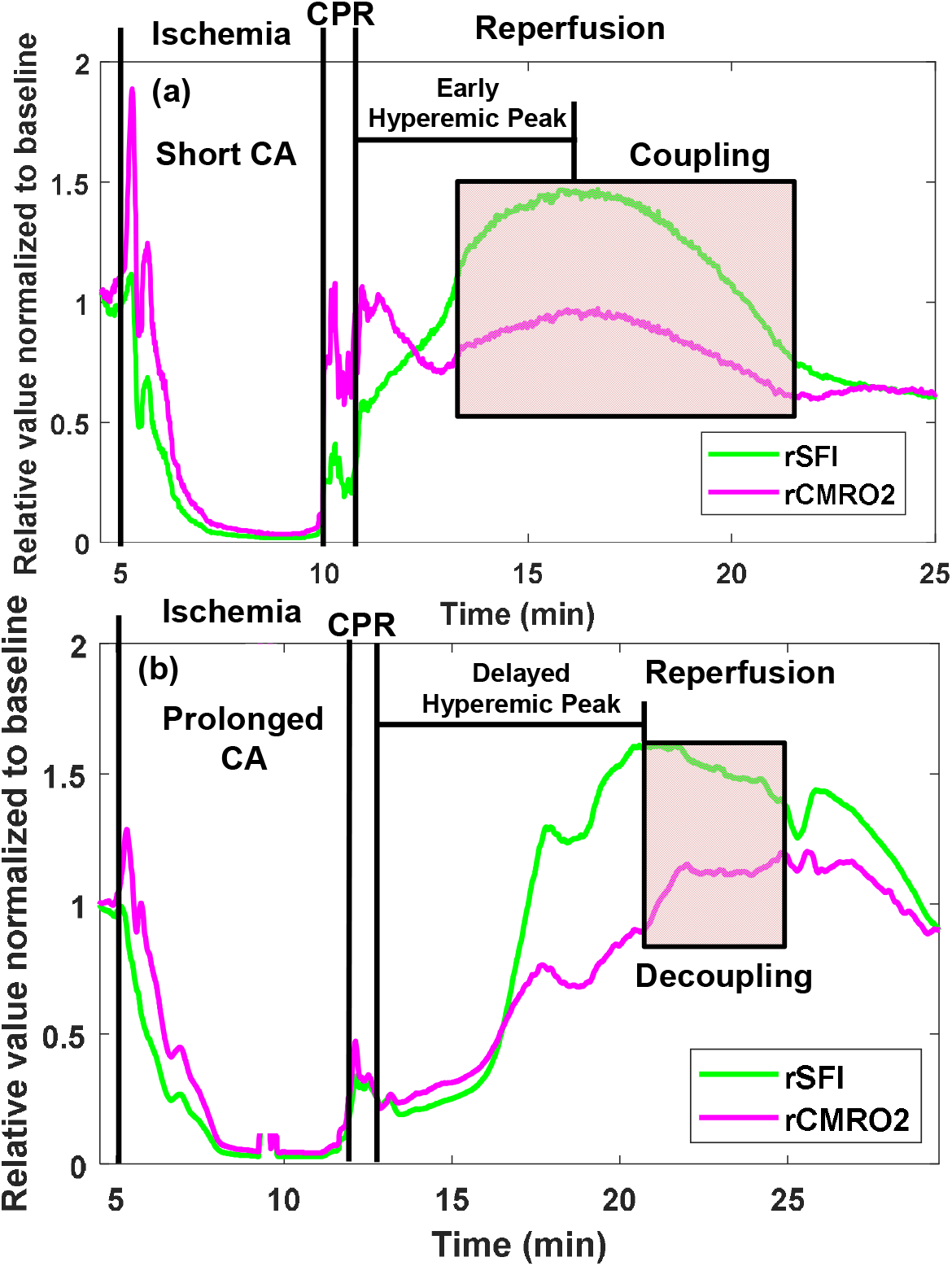
(a) A representative rat with short CA (5 min asphyxia) illustrates early, temporally-synchronous (coupled) recovery of CBF and CMRO_2_ in the post-ROSC period, (b) A representative rat with prolonged CA (7 min asphyxia) shows delayed recovery of CBF and CMRO_2_ in the post-ROSC period with periods of decoupling between CBF and CMRO_2_ temporal dynamics (shaded box).

### Flow-metabolism mismatch in the brain within the first minute post-CPR can assess CA duration and predict cerebral electrical recovery

Next, we focused on flow/metabolism mismatch, which we can measure by calculating the ratio of CBF and CMRO_2_. Fig. 5(a) shows the CBF/CMRO_2_ ratio during the first minute after resuscitation for five rats with shorter CA (5 min asphyxia; solid lines) and five rats with prolonged CA (7 min asphyxia; dashed lines). All data were normalized to 15 sec post-ROSC, so no pre-ROSC information was required. This figure shows that a threshold can be placed on the value of CBF/CMRO_2_ at 1 min post-resuscitation (vertical line at CBF/CMRO_2_ ~ 1 in Fig. 5(b)) to separate the rats with shorter CA from the rats with prolonged CA. This result suggests that within 1 min of resuscitation, the CBF/CMRO_2_ ratio can be used for assessment of the severity (duration) of CA, <u>*without any prior knowledge of the cardiac or hemodynamic history of the patient*</u>. This threshold makes sense physiologically because CBF/CMRO_2_ < 1 can be thought of as a marker of flow-metabolism mismatch (i.e. CBF is insufficient to meet metabolic demand). The capability of the CBF/CMRO_2_ ratio to assess CA severity vanished within 3 min of ROSC.

**Fig. 5.**
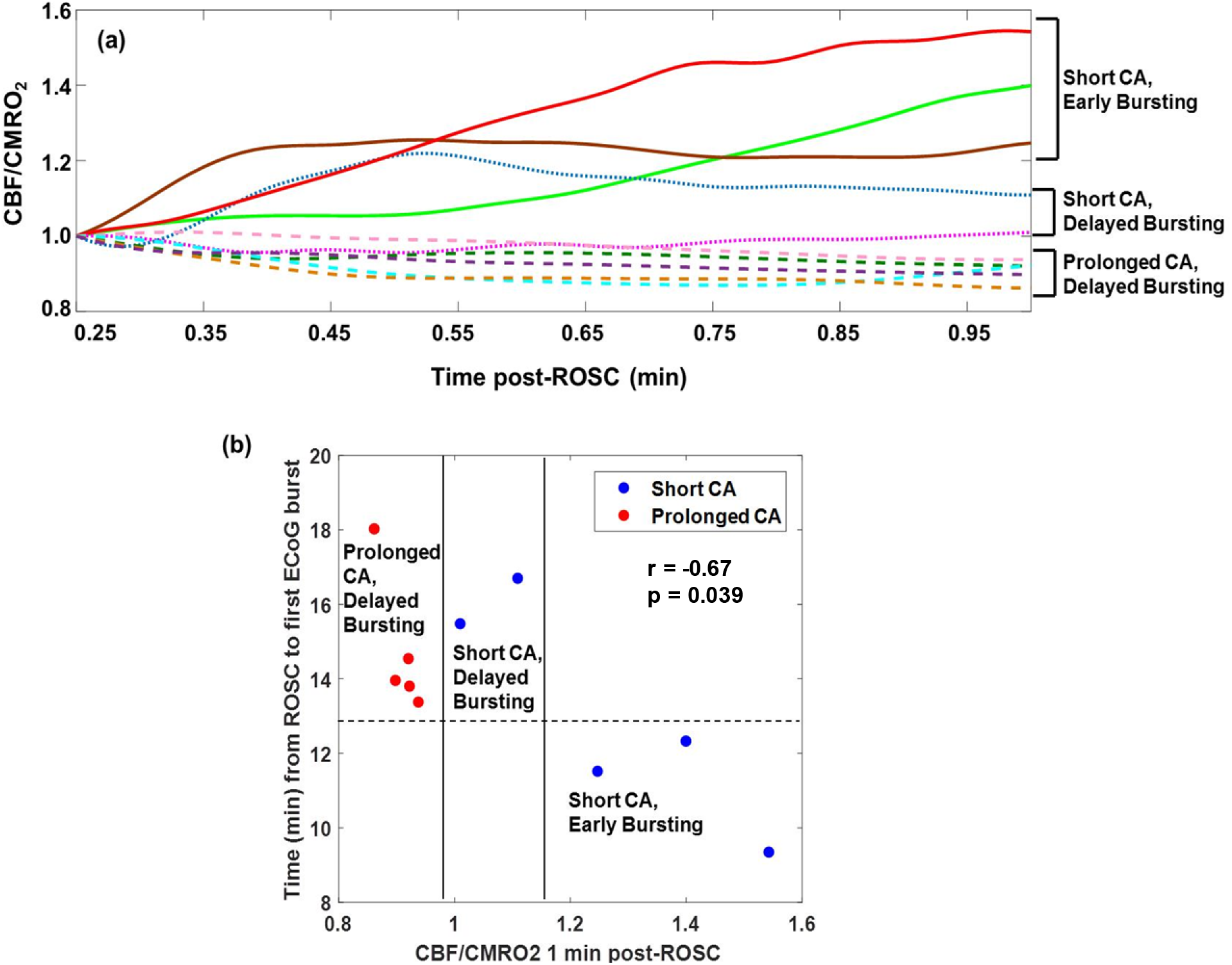
Ratio of CBF to CMRO_2_ (normalized to 15 sec post-ROSC) within the first minute post-ROSC can be used to retrospectively determine severity of CA and simultaneously provide a preliminary prediction of expected outcome. This ratio does not require any pre-ROSC information, making it well-suited for potential translation to emergency response and intensive care settings. (a) Using CBF/CMRO_2_ values, the time window of ~0.5-2 min post-ROSC is the most useful for CA severity assessment and prognosis. For instance, at 1 min post-ROSC, a clear separation is observed between rats that had prolonged CA (7 min asphyxia) and rats that had shorter CA (5 min asphyxia). Rats that had short CA but ended up experiencing delayed ECoG bursting have CBF/CMRO_2_ values that fall above those of the rats with prolonged CA but below the other rats with shorter CA. (b) CBF/CMRO_2_ 1 min post-ROSC correlates with time from ROSC to first ECoG burst (r = −0.67, p = 0.039 from Spearman correlation). A threshold of CBF/CMRO_2_ ~ 1 (dashed vertical line), suggestive of paired flow/metabolism coupling, can completely distinguish rats that had shorter duration of CA (5 min asphyxia) from prolonged CA duration (7 min asphyxia). A second threshold of CBF/CMRO_2_ ~1.2 completely distinguished rats that had good short-term cerebral electrical recovery (earlier ECoG bursting) from rats that had poor short-term cerebral electrical recovery (later ECoG bursting), regardless of CA duration. These correlations vanish within the first 3 min post-ROSC, suggesting the presence of a transient ultra-early window for assessing CA severity and predicting neurological recovery.

Furthermore, a second threshold can be placed at CBF/CMRO_2_ ~ 1.2 to differentiate between rats with poor short-term recovery (longer time to ECoG bursting) and those with good short-term recovery (shorter time to ECoG burst), independent of CA duration. Fig. 5(b) shows a scatter plot for the values of this ratio for all rats, plotted against the time to initial ECoG burst. The relationship between CBF/CMRO_2_ and initial ECoG burst was statistically significant using a Spearman correlation (r=-0.67, p=0.039). Overall, a higher flow/metabolism index at 1 min post-ROSC is associated with a shorter asphyxial CA period and a better neurological outcome as measured by faster ECoG bursting. The ECoG burst time prediction capability of the CBF/CMRO_2_ ratio also vanished within 3 min of ROSC.

To test the prognostic ability of the CBF/CMRO_2_ ratio, we created a predictive model by performing leave-one-out cross-validation (35) with linear fits to the points on the scatter plot of ECoG burst time vs. CBF/CMRO_2_ at 1 min post-ROSC. Based on this technique, the CBF/CMRO_2_ ratio was predictive of the initial burst time with 87% accuracy (Supplementary Table 1). Importantly, the CBF/CMRO_2_ mismatch ratio provided both CA severity assessment and recovery prognosis simultaneously within an ultra-early time window (~0.5-2.0 min post-ROSC).

To further analyze the impact of flow-metabolism mismatch immediately post-ROSC on early neurological recovery, we calculated additional indices related to the difference between CBF and CMRO_2_ and compared these indices to the time of the first EEG burst post-ROSC. Supplementary Figs. 1(a, b) show CBF and CMRO_2_, normalized to the corresponding value at 15 sec post-ROSC, for the first 5 min post-ROSC for representative rats with shorter CA (Supplementary Fig. 1(a)) and longer CA (Supplementary Fig. 1(b)). During this period, the relative CBF is higher than the relative CMRO_2_ for the rat with shorter CA, but the relative CMRO_2_ is higher than the relative CBF for the rat with the prolonged CA. These trends suggest that higher CBF in comparison to metabolic demand by the brain is associated with shorter CA duration. Supplementary Fig. 1(c) shows that the ratio of the areas under the CBF and CMRO_2_ curves from 0.25-3.00 min post-ROSC were different for shorter CA and prolonged CA.

### Rate of change of CBF within the first minute post-ROSC correlates with time to resumption of cerebral electrical activity

Fig. 6 shows that CBF alone, measured within the first minute post-ROSC and normalized to its value at 15 sec post-ROSC, can be employed to predict the time of initial ECoG burst, without knowledge of CMRO_2_. A linear regression (black line; r = −0.77, p = 0.01) was fit to the data in Fig. 6. Using a leave-one-out cross-validation technique, this metric predicted first ECoG burst to within 16% over the entire cohort of rats, including both shorter and prolonged asphyxia times. This result suggests that the lower the CBF after completion of CPR, the longer it will take for the brain’s electrical activity to resume. This finding is corroborated by our previously-reported result (9) that there is a certain threshold on total perfusion required for ECoG bursting to begin following ROSC. However, the result reported here suggests that knowledge of the total time-integrated perfusion is not required to predict ECoG bursting; only the change in CBF 30 sec post-ROSC relative to its value at 15 sec post-ROSC is required. It is important to note that CBF was not as accurate as the CBF/CMRO_2_ ratio for assessing CA duration in this time window.

**Fig. 6.**
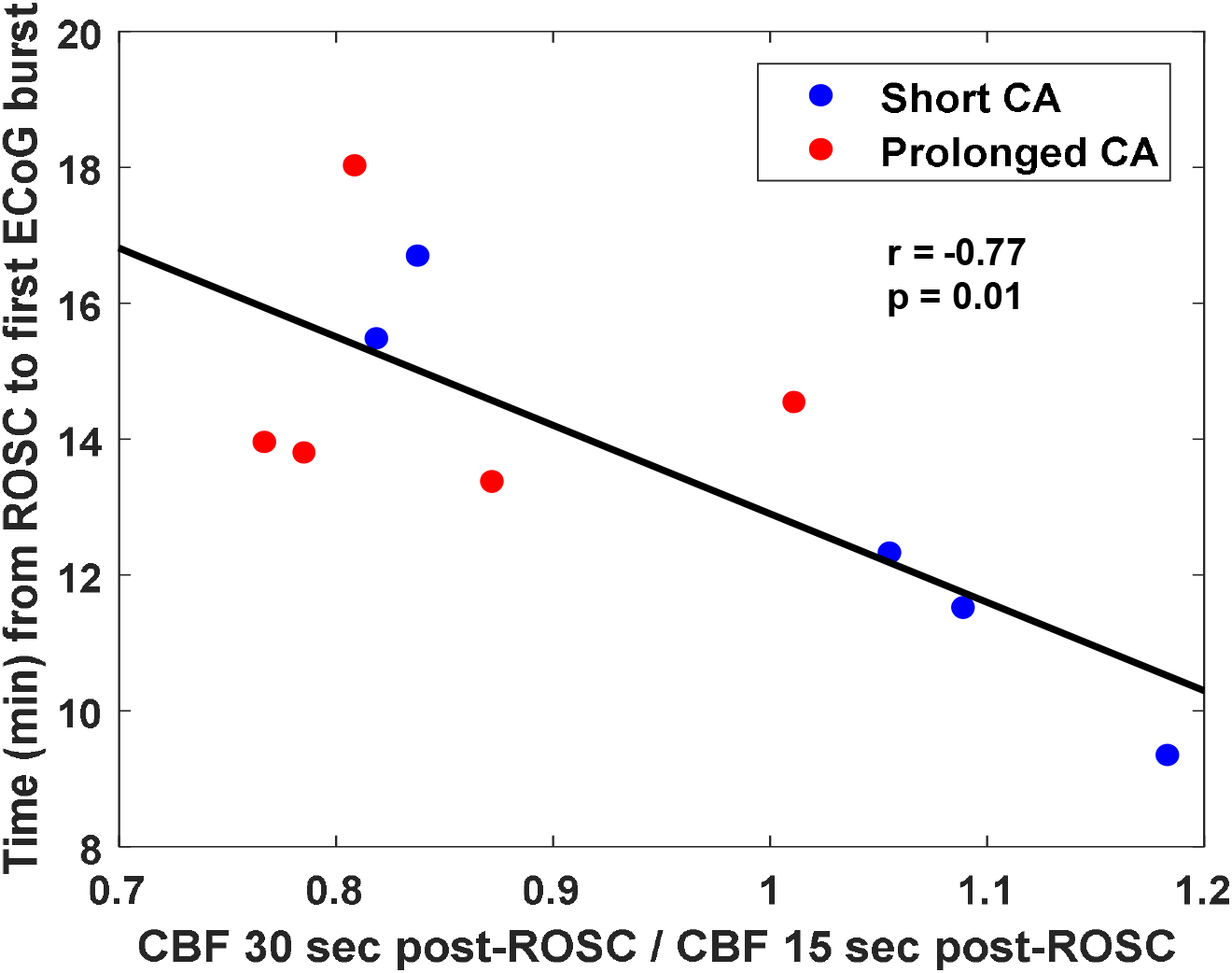
(a) CBF, measured at 30 sec post-ROSC and normalized to its value at 15 sec post-ROSC, is linearly related to ECoG burst time (black line; r = −0.77, p = 0.01 from Pearson correlation). This correlation vanished within 2 min after ROSC.

## Discussion

### Impaired autonomic regulation highlights importance of flow-metabolism metrics during a critical time window immediately post-resuscitation

In the healthy brain, autonomic regulation is intact, so a neural stimulus will trigger an appropriate increase in CBF, matched with the corresponding increase in metabolic demand (28). Typically, the CBF response will overshoot the increase in metabolism; this appears to be a normal physiological reaction designed to maintain a reserve supply of oxygen in case metabolic demand increases or the ambient oxygen level decreases (36). This type of system, demonstrated often in stimulus-evoked response experiments (34), is a classic example of optimal neurovascular coupling and intact autonomic regulation. However, after acute cerebral ischemia or other forms of brain trauma, cerebral autonomic regulation may be compromised, causing impaired neurovascular coupling and mismatches between CBF and metabolism (37). Therefore, it is critical to obtain better quantitative understanding of flow-metabolism mismatch immediately following these types of insults because CBF, blood pressure, oxygenation, or cortical electrical activity alone may not provide an accurate picture of brain function and neural dynamics during these critical time periods.

Here, we find that deviations of the CBF/CMRO_2_ ratio from unity *within the first minute post-ROSC* can assess CA severity (asphyxia duration) and predict cerebral electrical recovery (time to first ECoG burst) (Fig. 5). The CBF/CMRO_2_ ratio at 1 min post-ROSC is predictive of ECoG burst time with 87% accuracy (Supplementary Table 1). At 30 sec post-ROSC, using CBF alone (Fig. 6) can predict ECoG burst time with 86% accuracy. Note that these metrics do not require any pre-ROSC information.

Fig. 7 provides a timeline over which the CBF and CBF/CMRO_2_ metrics are significant for assessing CA severity and predicting short-term cerebral electrical recovery (ECoG burst time) post-ROSC. As seen in Fig. 7, CBF alone may provide an ultra-early (~0.5 min post-ROSC) prognostic marker. However, the CBF/CMRO_2_ ratio is required to perform severity assessment and recovery prognosis in tandem, and the critical window for simultaneously obtaining both of these pieces of information is ~1-2 min post-ROSC. The significance for all of these metrics vanishes within 3 min of ROSC, suggesting that the first ~3 min post-ROSC may provide a critical but transient window during which therapeutic maneuvers can be investigated for improving neurological outcome after CA.

**Fig. 7.**
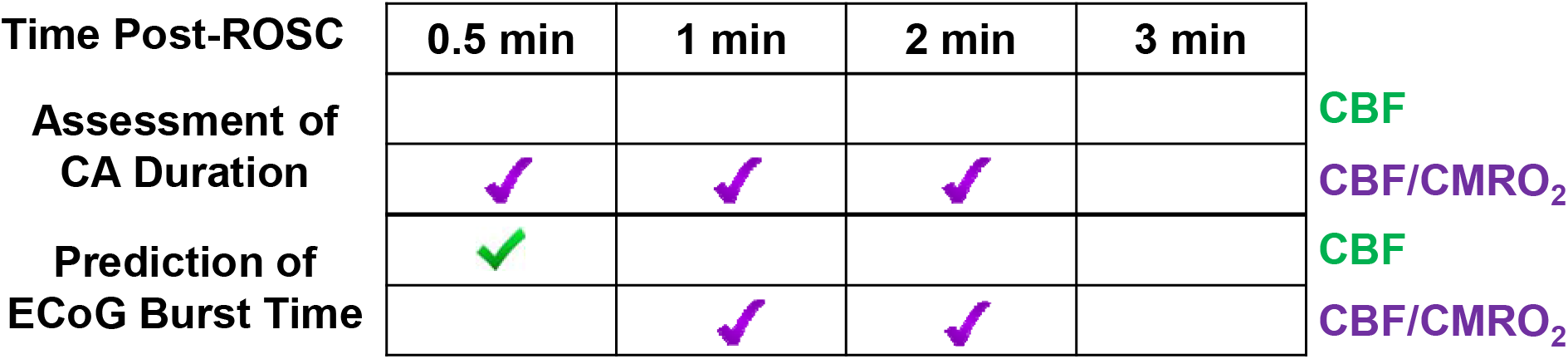
Post-ROSC timelines over which CBF alone (green checkmark) and CBF/CMRO_2_ (purple checkmark) are statistically significant for assessing CA duration and prognosticating resumption of ECoG bursting. Significance for assessing CA duration was determined with Wilcoxon rank-sum tests for distinguishing between 5 and 7 min asphyxial times. Significance for prognosis of ECoG burst time was assessed via Spearman correlations. Our findings support the existence of a transient, ultra-early critical window from ~0.5-2.0 min post-ROSC during which CA duration can be assessed and ECoG burst time can be predicted. CBF alone (at 0.5 min post-ROSC) can predict ECoG burst time, but the CBF/CMRO_2_ ratio is required to obtain simultaneous assessment of CA duration and prognosis of cerebral electrical recovery. All data were normalized to their values at 15 sec post-ROSC, so no pre-ROSC information was required for these calculations.

### Immediate flow-metabolism monitoring is critical for improving CA patient outcome post-CPR

CA patients typically suffer pronounced and prolonged brain damage due to cerebral ischemia. For patients who undergo out-of-hospital CA, 68% of fatalities are attributed primarily to ischemia-related brain injury (3), and fewer than 9% survive with “Good or Moderate Cerebral Performance” (defined as Cerebral Performance Category 1 or 2) (38, 39). With the exception of targeted temperature management, there are no widely-accepted clinical treatments to improve CA patient outcome. Also, developing diagnostic and prognostic tools to optimize blood pressure, oxygen, and carbon dioxide levels for these patients is an active area of investigation (40, 41). Recently, it has been suggested that increasing the mean arterial pressure immediately post-ROSC may mitigate flow-metabolism uncoupling by maximizing CBF (42). However, it is also known that too high of a CBF or oxygenation level during this critical period can potentially increase the risk of reperfusion injury and oxidative damage to mitochondria and neurons (5, 43). Therefore, there is an unmet clinical need for real-time quantitative monitoring of CBF and brain metabolism following CA, especially in the transient hyper-dynamic period immediately post-CPR. In the intensive care setting, brain function of CA patients is typically monitored with electroencephalography (EEG), and perfusion is usually assessed via peripheral blood pressure. As a result, the underlying mechanisms driving recovery of cerebral electrical activity following hypoxic-ischemic injury are not well-characterized, and measuring them could lead to improvements in patient care.

### Cerebral perfusion/metabolism mismatch can predict ischemic injury or reperfusion damage to prognosticate neurological recovery and inform treatment

We postulate that an optimal CBF range exists to promote cerebral recovery following an ischemic injury such as CA, but this range can vary between patients and becomes narrower after hypoxic-ischemic events (7). Furthermore, this optimal range is defined not by the CBF alone, but by the amount of perfusion relative to cerebral metabolism (44). Determining this optimal balance of flow-metabolism matching to allow optimal neurovascular coupling is especially critical during periods of cerebral autonomic dysregulation, which occurs after acute brain injury (including ischemic injury (37) and traumatic brain injury (45)). Therefore, measuring CBF and CMRO_2_ in tandem is crucial, and the CBF/CMRO_2_ ratio may be used to indicate ischemic damage (CBF/CMRO_2_ < 1) or excess perfusion (CBF/CMRO_2_ >>1).

In our model, we find that it is optimal to have a CBF/CMRO_2_ ratio that exceeds 1 in the first few minutes post-CPR; this ratio may need to be much greater than 1 to indicate reperfusion injury. By contrast, at 1 min post-ROSC, a CBF/CMRO_2_ ratio that is even slightly below 1 (or, in fact, slightly above 1) may indicate risk of ischemic injury, as animals with delayed ECoG bursting in our study had CBF/CMRO_2_ < 1.2 at this early time point. It is important to remember that the significant prognostic metrics in this report were all found at time points within ~3 min post-ROSC. After that time window ended, these metrics lost their prognostic significance (e.g., Fig. 5). The transient nature of this prognostic window, which may easily be overlooked, may be a potential explanation for why it is currently difficult for clinicians to determine the optimal blood pressure for post-CA patients in the intensive care unit. Specifically, peripheral blood pressure may be decoupled from CBF (9), measurements of CBF are not typically combined with cerebral oximetry (46), and there is often a significant time delay between ROSC and measurements of cerebral perfusion. Continuously monitoring CBF and CMRO_2_ immediately post-ROSC may provide real-time feedback to clinicians to optimize treatment and improve cerebral recovery for CA patients.

### Measuring flow-metabolism mismatch can provide early assessment of CA severity/duration

In addition to prognosis of cerebral electrical recovery, the CBF/CMRO_2_ ratio at 1 min post-ROSC provided complete distinction between rats that had undergone mild CA (5 min asphyxia) and rats that experienced more severe CA (7 min asphyxia) (Fig. 5). Specifically, rats that experienced more severe CA had lower CBF/CMRO_2_ ratios (suggesting ischemic injury) at this time point than rats with milder CA. Therefore, quantifying the flow-metabolism mismatch post-resuscitation may also provide retrospective assessment of severity/duration of ischemia. Obtaining this assessment can be transformational in clinical management and prognostication of post-CA patients because true “down-time” (hypoxic-ischemic duration) is often not known when first responders arrive on the scene. Furthermore, for the two rats that underwent mild CA but exhibited delayed resumption of ECoG activity, the CBF/CMRO_2_ ratio was closer to those of the animals with more severe CA. This result suggests that quantifying cerebral perfusion-metabolism mismatch can potentially provide finer stratification of CA severity assessment and recovery prognosis across multiple subgroups of CA/CPR patients.

### Limitations of this study

Limitations exist in our methodology. Our sample size was small (N=10); in future work we will further validate these findings in a larger cohort. Additionally, the experiments were not survivable, because we used an open craniectomy procedure that exposed the brain. As a result, we were unable to obtain behavioral data or quantitative ECoG parameters to assess longer-term neurological recovery. Furthermore, the craniectomy can cause changes in intracranial pressure (potentially affecting blood pressure and blood flow), and exposing the brain to air may alter cerebral oxygenation and hydration. Future work will involve the use of a cranial window or thinned skull preparation (47). The cranial window will also enable longer-term monitoring of animals out to several weeks post-CPR to assess longer term neurological outcome and behavioral endpoints. Finally, our imaging technologies are primarily sampling the cortex; incorporation of fiber-probe based methods can potentially be incorporated into future studies to continuously interrogate regions of the brain that are deeper beneath the surface (48).

### Future work

Future studies should aim to validate the clinical translational potential of the aforementioned flow-metabolism mismatch metrics in larger animals (i.e. pigs) and humans. Applying non-invasive near-infrared spectroscopy (NIRS) and diffuse correlation spectroscopy (DCS) to measure CBF and CMRO_2_ immediately post-ROSC in CA patients will be of high value. In the preclinical model, it will be of interest to modify CBF within, and outside of, the critical window of ~0-3 min post-ROSC and examine the effects of these interventions on short-term neurological recovery (e.g., ECoG bursting) and longer-term outcome (e.g., neurological arousal, behavioral, and cognitive tests 24 hrs to 1 week post-ROSC). Furthermore, it will be valuable to rapidly modify CMRO_2_ via hypothermia or other approaches at different time points within or outside of this critical window to examine its effect on short-term recovery and longer-term outcome (49–52). These studies will be important for obtaining a better understanding of the causal relationship between flow-metabolism mismatch, neurovascular coupling, and neurological recovery following CA.

## Conclusion

In this report, we quantify the highly-dynamic relationship between CBF and brain metabolism (CMRO_2_) in a preclinical model of CA and CPR. We observe different degrees of coupling between CBF and CMRO_2_ in different temporal windows over the first ~20 min following CPR. We calculated the degree of flow-metabolism mismatch by using the metric CBF/CMRO_2_. This mismatch was significant for assessing CA severity and prognostically significant (correlating with time to initial ECoG burst) *within the first minute post-ROSC*. However, the statistical significance of these correlations vanishes within ~3 min post-ROSC, suggesting the presence of a transient, critical time window during which continuous monitoring of CBF and CMRO_2_ may be crucial for assessment of severity of hypoxic brain injury, prognostication, and optimizing potential treatments for CA patients.

## Acknowledgements

This work is supported by the Arnold and Mabel Beckman Foundation, the United States National Institutes of Health (P41EB015890), the National Science Foundation Graduate Research Fellowship Program (DGE-1321846, to C.C.), the National Center for Research Resources and National Center for Advancing Translational Sciences, National Institutes of Health (through the following grants: R21EB024793 to Y.A., TL1TR001415-01 to R.H.W., 5KL2TR000147 to Y.A., a pilot grant to Y.A., all via UL1 TR001414, and a CTSA pilot grant to Y.A. via UL1 TR000153), and the Roneet Carmell Memorial Endowment Fund to Y.A.. The content is solely the responsibility of the authors and does not necessarily represent the official views of the NIH.

## Contributions

R. H. W. and C. C. performed data collection and analysis and manuscript writing; M. T. performed instrumentation development; A. B., N. M., and J. A. performed animal preparation, monitoring, and data collection; B. J. T. and B. C. provided scientific direction and edited the manuscript; Y. A. provided scientific direction, organization of the study, manuscript writing/editing, and resuscitation and monitoring of the animals in this study.

---

**Supplementary Fig. 1.**
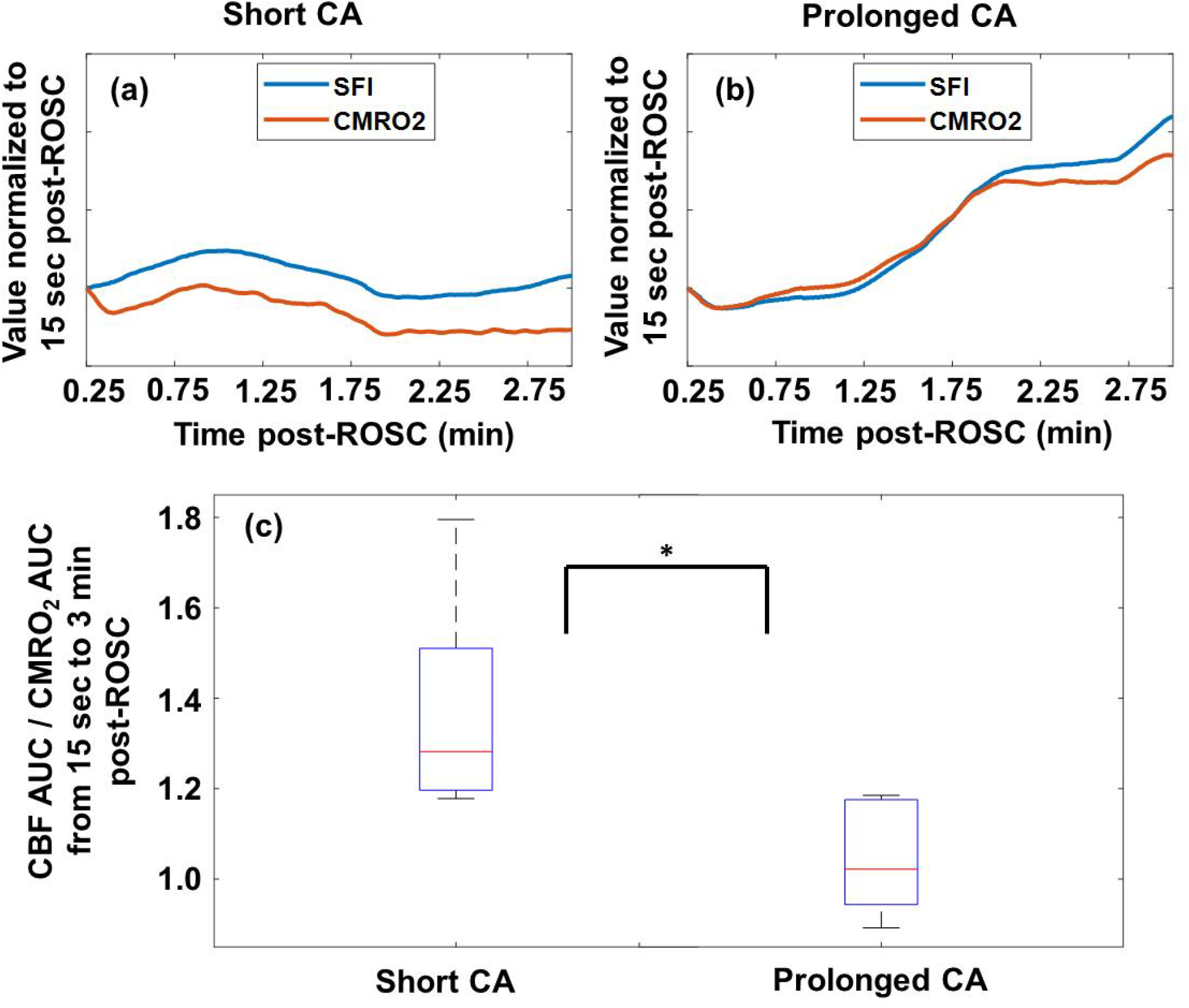
CBF exceeds CMRO_2_ during first 5 min post-ROSC for a representative rat with short (5 min, (a)) CA, but not for prolonged (7 min, (b)) CA. (c) The ratio of the areas under the CBF and CMRO_2_ curves (AUC) from 15 sec to 3 min post-ROSC is significant (*, p < 0.02 from Wilcoxon rank-sum test) for distinguishing between rats that underwent short CA and those with prolonged CA. Prior to the AUC calculation, the CBF and CMRO_2_ were normalized to their values at 15 sec post-ROSC. No pre-ROSC information was required for this calculation.

**Supplementary Table 1.**
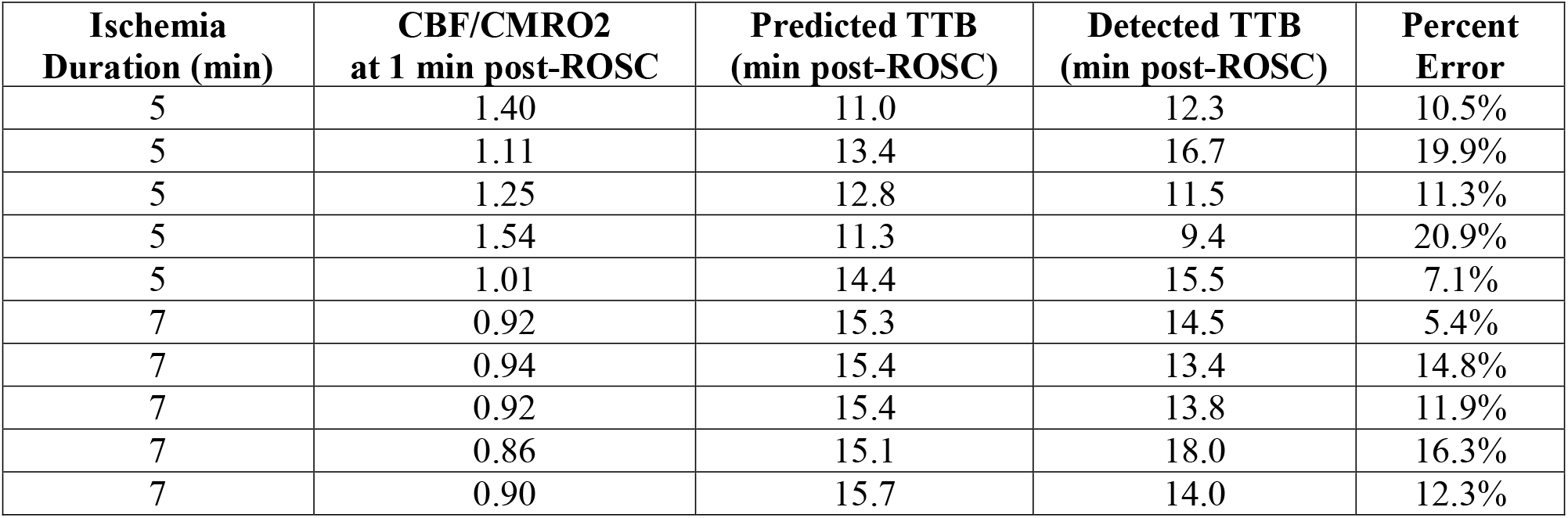
The ratio between CBF and CMRO_2_ (CBF/CMRO_2_; column 2) 1 min post-ROSC can be input into a linear regression model to predict time to initial ECoG burst (TTB, detected using our previously-published algorithm (9)). Using a leave-one-out cross-validation technique, the mean percent error for predicting TTB was 13% over the full cohort of rats in this study, and the error did not exceed 21% for any of the rats. Prior to this calculation, CBF and CMRO_2_ were normalized to their respective values at 15 sec post-ROSC. The method did not require any pre-ROSC information.

| **Ischemia Duration (min)** | **CBF/CMRO2 at 1 min post-ROSC** | **Predicted TTB (min post-ROSC)** | **Detected TTB (min post-ROSC)** | **Percent Error** |
| --- | --- | --- | --- | --- |
| 5 | 1.40 | 11.0 | 12.3 | 10.5% |
| 5 | 1.11 | 13.4 | 16.7 | 19.9% |
| 5 | 1.25 | 12.8 | 11.5 | 11.3% |
| 5 | 1.54 | 11.3 | 9.4 | 20.9% |
| 5 | 1.01 | 14.4 | 15.5 | 7.1% |
| 7 | 0.92 | 15.3 | 14.5 | 5.4% |
| 7 | 0.94 | 15.4 | 13.4 | 14.8% |
| 7 | 0.92 | 15.4 | 13.8 | 11.9% |
| 7 | 0.86 | 15.1 | 18.0 | 16.3% |
| 7 | 0.90 | 15.7 | 14.0 | 12.3% |

